# Evidence for a sensorimotor prediction process in action observation

**DOI:** 10.1101/2023.02.04.527111

**Authors:** Pauline M. Hilt, Nicolas Gueugneau, Luciano Fadiga, Thierry Pozzo, Charalambos Papaxanthis

## Abstract

The mirror neurons network in the human brain is activated both during the observation of action and the execution of the same action, facilitating thus the transformation of visual information into motor representations, to understand the actions and intentions of others. How this transformation takes place, however, is still under debate. One prevailing theory, *direct matching*, assumes a direct correspondence between the visual information of the actor’s movement and the activation of the motor representations in the observer’s motor cortex that would produce the same movement. Alternatively, the *predictive coding* theory postulates that, during action observation, motor predictions (e.g., position, velocity) are generated and compared to the visual information of the actor’s movement. Here, we experimentally interrogate these two hypotheses during a locomotion task. The motor prediction process was assessed by measuring the timing of imagined movements: the participants had to imagine walking, forward or backward, for 9 m (linear path). Action observation was assessed by measuring time estimation in an inference locomotor task (the same 9 m linear path): after perceiving an actor walking forward or backward for 3 m, the vision of the observer was occluded and he/she had to estimate when the actor would reach the end of the 9 m path. We manipulated the timing processes during the two tasks by creating sensory illusions via peripheral mechanical muscle vibration on leg muscles, which has provided consistent results in the literature (acceleration of forward and deceleration of backward locomotion). We found that sensory illusions specifically affected the timing processes of both locomotion inference and mental locomotion, suggesting the involvement of sensorimotor predictions, common to both tasks. These findings seem to support the predictive coding hypothesis.

## Introduction

The observation of Olympic gymnasts performing complex and elegant movement combinations is always a fascinating and pleasant moment. In everyday life, perceiving and understanding the actions of others is essential for social communication and learning. In the human brain, a similar neural network is activated both during the observation of action and the execution of the same action by the observer. This neural network, named *‘mirror neurons network’*, due to its particular properties, includes the motor areas (Ferroni et al., 2021; Vigneswaran et al., 2013; Albertini et al., 2021; Papadourakis & Raos, 2019), the parietal areas (Bonini et al., 2010; Lanzilotto et al., 2019, 2020), the prefrontal cortex (Simone et al., 2017), the basal ganglia, and the cerebellum (Errante & Fogassi, 2020). Note that *mirror neurons* have originally been discovered in area F5 of the monkey premotor cortex (di Pellegrino et al., 1992; Gallese et al., 1996; Rizzolatti et al., 1996). The involvement of a similar neural network for action observation and execution may be the neural basis of a functional process allowing humans to transform visual information into motor representations and thus to naturally understand the actions and intentions of others (Gallese et al., 2004, 2006; Jeannerod, 2001; Rizzolatti et al., 2001; Rizzolatti & Craighero, 2004).

How this transformation operates is, however, still under debate. According to the *direct matching* hypothesis, there is a direct correspondence between the visual information extracted from the actor’s movement and the activation of the motor representations in the observer’s motor cortex that would produce the same movement. Neurophysiological evidence suggests that corticospinal excitability in the observer’s motor system faithfully replicates the spatiotemporal sequence of motor commands implemented by the actor (Fadiga et al., 1995; for a review see: Naish et al., 2014). For example, observation of index and little finger abductions selectively increases the corticospinal excitability from the first dorsal interosseous and abductor digiti minimi muscles, responsible for the index and little finger movements, respectively (Catmur et al., 2011).

Although the *direct matching* hypothesis is very seductive due to its simplicity and could be operative for simple mono-articular movements, its effectiveness for more complex movements is more problematic. This is because there is no unique solution allowing someone to deduce from the observation of movement kinematics (here, the visual information of the actor’s movement) the appropriate motor commands to produce this same movement (here, the activation of the motor representation in the observers’ motor action); the well-known inverse problem. Indeed, for the production of multi-joint movements, due to the abundance of degrees of freedom, different joint configurations, as well as spatiotemporal patterns of muscle activity, can equally be used (Bernstein, 1967). Moreover, the *direct matching* hypothesis does not explain how an observer could learn/improve her/his motor repertoire by observing novel actions for which his/her motor representations are not yet been consolidated. There is, however, extensive literature showing the effectiveness of action observation in motor learning (Rizzolatti et al., 2021; Rizzolatti & Sinigaglia, 2016).

An alternative hypothesis, the *predictive coding*, based on the concept of motor prediction, could provide a solution to the inverse problem during action observation. Theoretically, when an individual prepares a movement, sensorimotor prediction is generated by internal forward models, which are neural networks that mimic the causal flow of the physical process by predicting the future sensorimotor state (e.g., position, velocity) given the efferent copy of the motor command and the current state (Wolpert & Flanagan, 2001). A self-supervised learning process minimizes prediction errors, namely the errors between the output from the forward model and the sensory outcome (sensory feedback) of the motor command. This process is vital for sensorimotor learning and adaptation (Wolpert et al., 2011). In the case of action observation, one could assume that visual information of the actor’s movement activates the observer’s motor representations, which in turn generate an internal copy of motor commands that are used from the internal forward model to make state predictions (e.g., position, velocity). These predictions could be compared to the visual information of the actor’s movement. If a mismatch exists, namely an error signal, it can be used to drive the controller to generate appropriate motor commands so that future prediction fits with the actor’s kinematics. Although the *predictive* hypothesis is more complex than the *direct matching* one, because it necessitates more operations implying further neural circuits, it has the advantage to provide a unique solution (from motor commands to kinematics, a forward process). While theoretical papers have argued in favor of the *predictive coding* hypothesis (Friston et al., 2011a; Keysers & Gazzola, 2014; Keysers & Perrett, 2004; Kilner & Frith, 2008), empirical data remains scarce. Few neurophysiological studies suggest that the motor system computes the difference between expected and observed action. A recent TMS study showed that during observation of complex multi-joint actions, the primary motor cortex activation reflects differences, rather than similarities, between the perceived movements and the observer’s motor patterns (Hilt et al., 2020).

Here, we tried to distinguish between the *predictive coding* and the *direct matching* hypothesis during action observation. We used a locomotor inference task: the participants (observers) watched an actor walking and had to evaluate the moment at which he/she would reach the end of the path. The observers had full vision only of the first 3 m of the actor’s walk, during the last 6m their vision was occluded. To measure motor prediction, we used motor imagery, in which participants performed the same 9 m locomotor path mentally. Forward models are believed to be involved in mental movement simulation (Miall & Wolpert, 1996; Wolpert & Flanagan, 2001) and temporal features of mental movements emerge from sensorimotor predictions (Papaxanthis et al., 2012; Sirigu et al., 1996). Finally, we manipulated action observation (locomotion inference) and prediction (mental locomotion) processes, by creating sensory illusions via peripheral mechanical muscle vibration on leg muscles, which has provided consistent results in the literature (Ivanenko et al., 2000). As sensory illusions did affect sensorimotor predictions during motor imagery (Kitada et al., 2002; Naito et al., 2002a; Thyrion & Roll, 2009), by creating persistent directional effects, they constituted an excellent way to test whether they could also affect sensorimotor predictions during action observation (*predictive coding* hypothesis). The absence of such a specific effect during action observation would validate the direct matching hypothesis.

## Results

In the following experiments, we bilaterally vibrated the tibialis anterior and hamstring muscles (see Figure 1). All the participants, without having any vibratory experience in the past, were sensible to the vibratory stimulation and presented illusory and postural reaction effects, similar to those described in the literature (Ivanenko et al., 2000). Precisely, tibialis anterior muscle vibration (85Hz) induced a prominent forward body tilt (on average trunk inclination 6.9 ± 2.6°) and an illusory perception of backward body tilt (verbally reported by all the participants). Hamstring muscle vibration (85Hz) induced backward inclination of the trunk (on average −7.3 ± 3.8°) and forward inclination of the shank (on average 9.4 ± 3.5°) and an illusory forward leg flexion relative to the trunk (verbally reported by all the participants).

**Figure 1.**
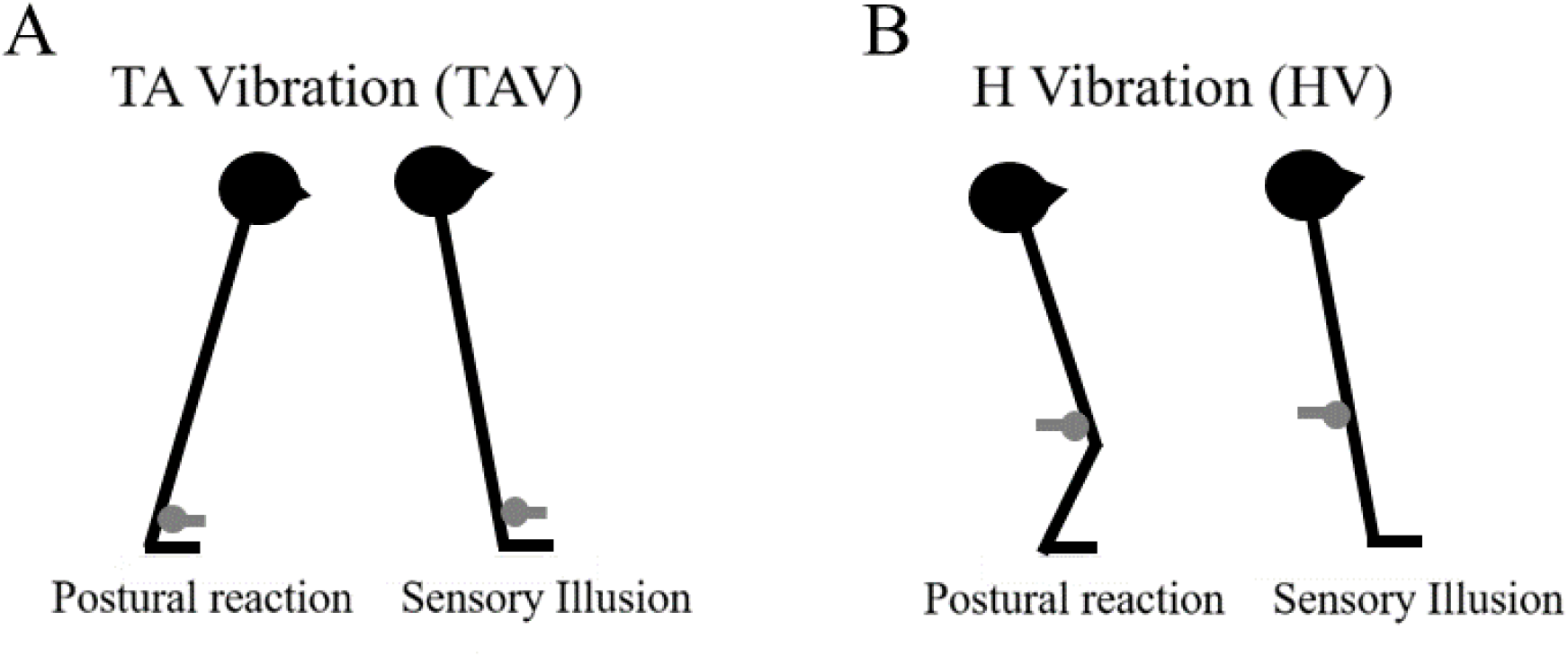
Muscle Vibration. The tibialis anterior (A) and hamstring (B) muscles were bilaterally stimulated by two mechanical vibrators at 85 Hz. The vibrators were fixed on the tibialis anterior (~4 cm above the ankle joint) and the hamstring distal (~3 cm above the knee joint) tendons. Qualitative illustrations of the postural and sensory illusions under muscle vibration are also illustrated. Tibialis anterior muscle vibration (A) induced a forward body tilt and an illusory perception of backward body tilt. Hamstring muscle vibration induced a backward inclination of the trunk and a forward inclination of the shank and an illusory forward leg flexion relative to the trunk (B).

### Experiment 1: Imagined locomotion under muscle vibration

In the first experiment (Figure 2A), we tested motor prediction through the motor imagery paradigm. We aimed to first understand how muscle vibration could influence motor prediction, precisely the duration of imagined locomotion, before looking for similar effects on action observation. In addition, as we wanted participants to be free of any experience regarding the effects of muscle vibration on actual locomotion, we included only imagined movements in this experiment.

**Figure 2.**
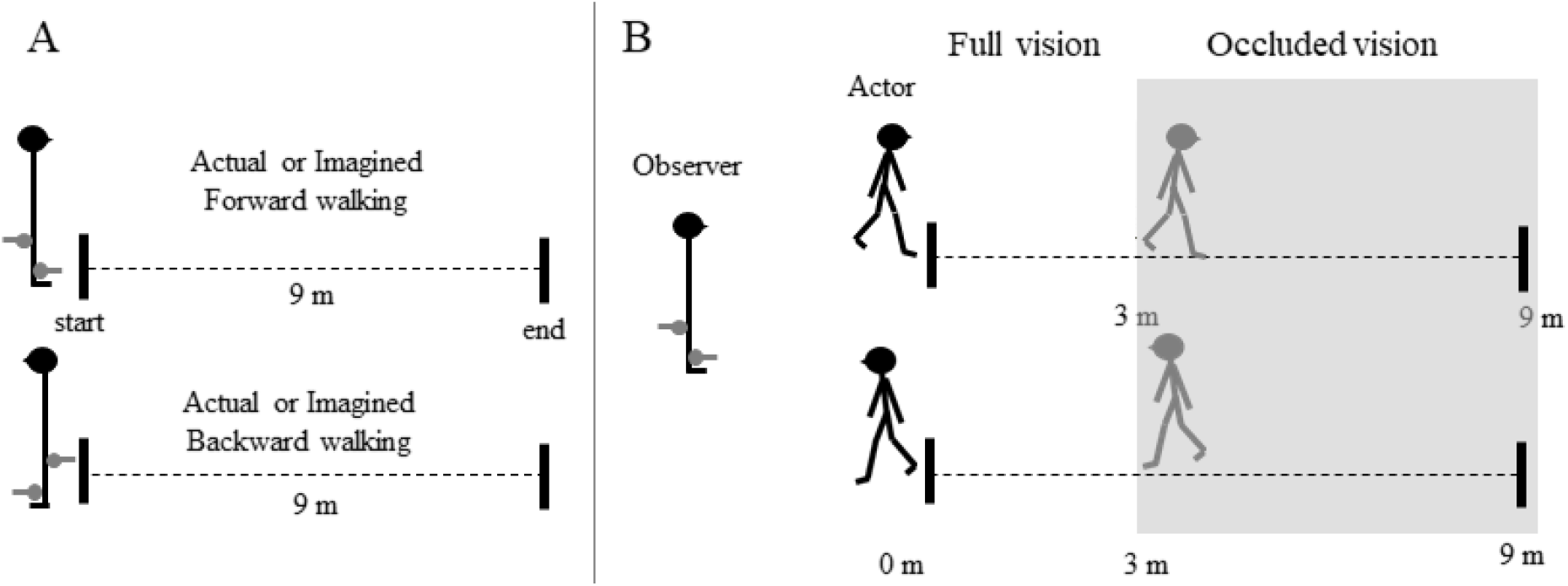
Experimental procedures. (A) Imagined and actual locomotion. Participants mentally (Experiment 1) and actually (Experiment 2) walked forward and backward along a 9m linear path at a natural speed with full vision and with or without vibration of the tibialis anterior and hamstring muscles. (B) Locomotion inference (Experiment 3). The participant (observer) watched a human actor walking along the 9m path (forward, top; backward, bottom) at a natural speed with or without vibration of the tibialis anterior and hamstring muscles. When the actor reached 1/3 of the path (3m) the observer’s vision was occluded thanks to crystals goggles until the end of the trial. The observer thus had to estimate when the actor would reach the end of the path.

Nine volunteers had to imagine walking at a natural speed along a path of 9 m in a forward or backward direction, with eyes open, to facilitate the visualization of the path, and under three vibration conditions: No-Vibration (NoV), bilateral vibration of the tibialis anterior muscle (TAV), and bilateral vibration of the hamstring muscle (HV). Two black lines on the floor materialized the initial and the final position of the path. The imagined duration of each trial (12 in each condition), was recorded using an electronic chronometer. Participants started the stopwatch when they mentally started to walk and stopped it when they mentally reached the end line. All participants reported vivid kinesthetic sensations in both walking directions and under all conditions (on average= 5.53 ± 0.96, on a 7-points Likert scale).

The average duration (± SD) of imagined locomotion is illustrated in Figure 3A. We obtained a significant interaction between *vibration* and *direction* (F_(2,16)_=112.6, p<0.0001, η^2^ = 0.94). Forward-imagined locomotion was faster than backward (for all conditions, p<0.001). Tibialis anterior muscle vibration did not affect imagined walking speed (p>0.1, for both forward and backward locomotion). In contrast, hamstring muscle vibration accelerated imagined forward locomotion (p<0.001; on average −8.25%; individual range from −4.6% to - 13.1%) and decelerated imagined backward locomotion (p<0.001; on average +7.59%; individual range from +5.7% to +9.8%). Walking imagined durations were stable through repetition (see supplemental Figure 1A). To test adaptation effects within the 12 trials of each experimental condition, we averaged the first three trials (1-3) and the last three trials (10-12) for each participant and compared them through two-tailed t-tests for dependent samples. The statistical analysis did not show any significant effect (for all conditions, t<1 and p>0.1).

**Figure 3.**
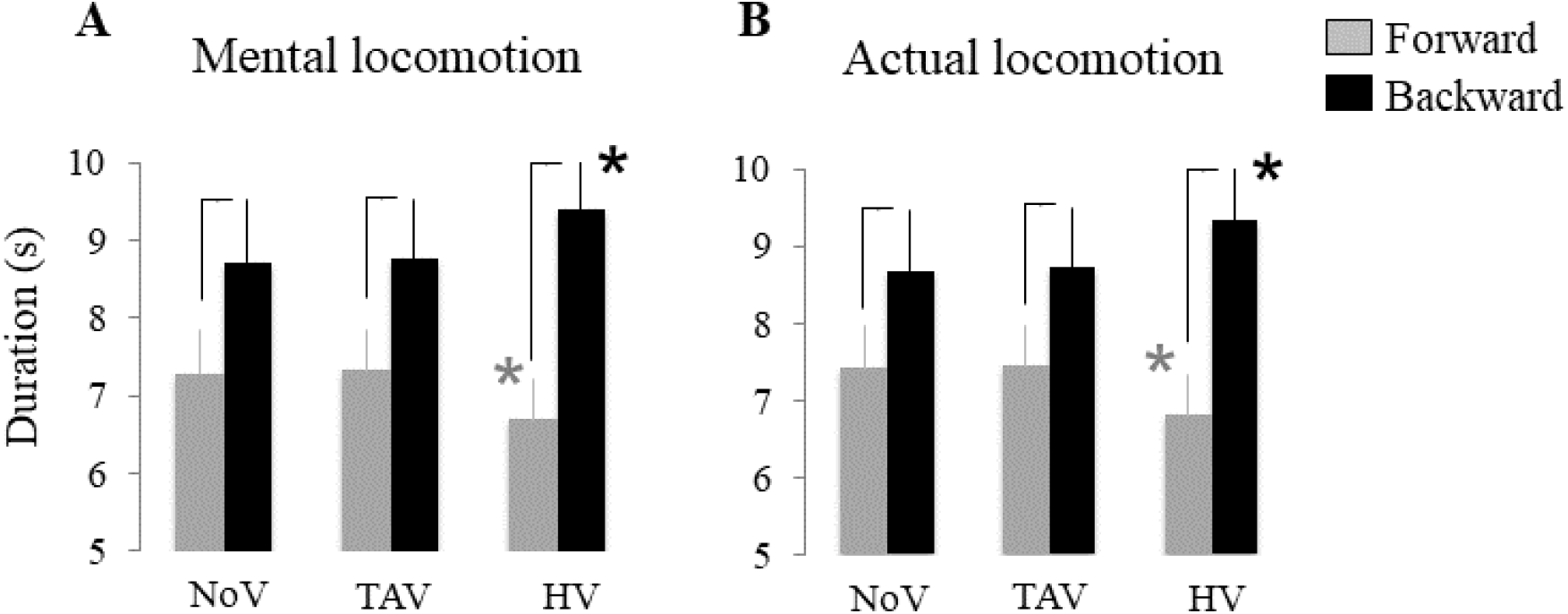
Mean durations (+ SD) of forward (grey) and backward (black) locomotion in the mental (A, Experiment 1) and actual (B, Experiment 2) locomotor tasks for each vibration condition. Black lines indicate significant differences between the walking directions. Grey and black stars indicate significant differences between the HV condition and the two others vibration conditions (NoV and TAV) within each direction of locomotion (forward and backward). NoV, no vibration; TAV, tibialis anterior muscle vibration; HV, hamstring muscle vibration.

Vibration effects on imagined locomotion were specific to the stimulation frequency (see supplemental Table 1). In the control experiment 1a, in which five participants of Experiment 1 realized the same protocol, the vibration of the hamstring muscle at 40 Hz did not alter forward (Permutation test; t=0.03, P=0.96) or backward (Permutation test; t=0.04, P=0.98) imagined walking speed.

Additionally, the main findings of Experiment 1 cannot be attributed to the postural effects elicited by muscle vibration (see Figure 1). Indeed, we did not observe any effect on the imagined walking speed (TAV-posture vs natural erect posture: Permutation test; t=-0.12, P=0.86; H-posture vs natural erect posture: Permutation test; t=0.04, P=0.96) when five participants of the Experiment 1 were asked to voluntarily adopt the postures elicited by tibialis anterior and hamstring muscles vibration during mental forward locomotion without vibration (control Experiment 1b, see supplemental Table 1).

### Experiment 2: Actual locomotion under muscle vibration

In the first experiment, we found that hamstring muscle vibration specifically affected the temporal aspects of imagined locomotion. In this second experiment, we tested whether similar effects were present during actual locomotion. The same participants were requested, at least one week later, to walk at a natural speed, with full vision, forward or backward, along the 9 m linear path, under the same vibration conditions (Figure 2B). Participants accomplished 12 trials for each experimental condition and walking duration was recorded using an electronic chronometer. Participants started the stopwatch when they started to walk and stopped it when they reached the end line.

Average walking durations (± SD) are illustrated in Figure 3B. ANOVA revealed a significant interaction between *vibration* and *direction* (F_(2,16)_=104.1, p<0.0001, η^2^ =0.93). Forward locomotion was faster than backward locomotion (for all conditions, p<0.001). While vibration of the tibialis anterior muscle did not affect walking speed (p>0.1, for both forward and backward locomotion), the vibration of the hamstring muscle accelerated forward locomotion (p<0.001; on average −8.72%; individual values ranged from −5.2% to −13.5%) and decelerated backward locomotion (p<0.001; on average +7.71%; individual values ranged from +6.3% to +9.9%). Walking durations under all vibration conditions were highly persistent (see supplemental Figure 1B) as a trial-by-trial (n=12) analysis did not reveal any adaptation through repetition (for all conditions, t<1 and p>0.1). These findings corroborate those of previous studies, which reported specific effects of leg muscle vibration on walking speed (Ivanenko et al. 2000).

We also verified in five of the previous participants (control experiment 2) that vibration of the hamstring muscle at 40 Hz did not alter forward (Permutation test; t=0.07, P=0.91) or backward (Permutation test; t=0.11, P=0.85) walking speed (see supplemental Table 1).

### Comparison between mental and actual locomotion

Hamstring muscle vibration induced almost similar effects on actual and imagined locomotion duration. Indeed, the statistical comparison of the vibratory effects, assessed by the ratio [(HV – NoV) / NoV*100)] and calculated for actual and imagined locomotion separately, did not show any effect neither for the forward (t=-1.85, p=0.10) nor the backward locomotion (t=0.55, p=0.59; Figure 4A). We found strong correlations between actual and imagined durations under hamstring muscle vibration for the forward (r=0.97, p<0.0001, t=11.3; Figure 4B) and backward (r=0.92, p<0.0001, t=6.35; Figure 4C) locomotion; the greater the effect of hamstring muscle vibration on actual locomotion, the greater its effects on imagined locomotion.

**Figure 4:**
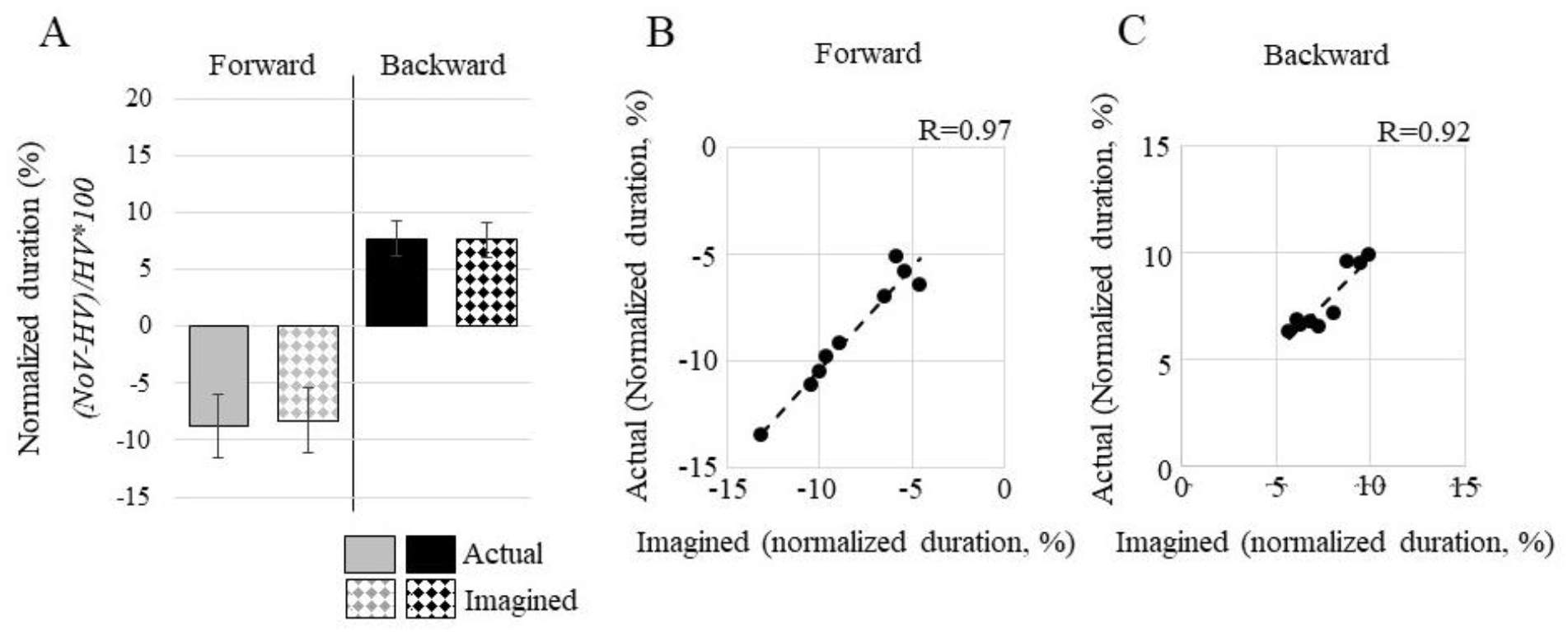
(A) Effects of hamstring muscle vibration (normalized by no vibration, NoV), on actual and imagined locomotion for forward (grey) and backward (black) walking. Correlations between actual (B) and imagined (C) durations under hamstring muscle vibration for the forward (r=0.97, p<0.0001, t=11.3) and backward (r=0.92, p<0.0001, t=6.35) locomotion; the greater the effect of hamstring muscle vibration on actual locomotion, the greater its effects on imagined locomotion.

### Experiment 3: Locomotion inference under muscle vibration

In the previous experiments, we found that the timing of imagined movements is directionally affected by muscle vibration, suggesting that motor imagery involves a sensory prediction process (see also Kitada et al., 2002; Naito et al., 2002a). The same findings observed during actual locomotion exclude the possibility that muscle vibration acted as an extra load that just perturbed speed locomotion.

Given these results, what could someone expect during action observation? To answer this question, we asked a new group of nine volunteers (4 females, 5 males, mean age: 25.2 ± 2.1 years) to participate in a locomotion inference task under muscle vibration. Each participant (hereafter ‘observer’) had to estimate the duration needed by another human (hereafter ‘actor’) to walk along the same 9 m linear path (Figure 2C). The actor walked forward or backward, with full vision, at natural velocity, without muscle vibration. A preliminary practice allowed the actor to adapt its speed to the natural speed of each observer with small variability. The observer stood upright in front of the actor at a distance of 2 m. The observer could fully perceive the actor’s locomotion for 1/3 of the path (i.e., 3 m). For the rest of the path (6 m, 2/3), the vision of the observer was occluded by crystal goggles with a wireless obstruction system (*Plato System*, 2 ms resolution). The actor’s walking duration and the corresponding temporal estimation by the observer were recorded using two electronic chronometers manipulated by the actor and the observer and synchronized via a computer program (see more details in the Method section). The observer realized the task (12 trials per condition) under the same vibration conditions (NoV, TAV, and HV, Figure 1) as in the previous experiments.

Based on the results of the previous experiments and the theoretical assumptions of the *predictive coding* and the *direct matching* hypothesis, we formulated the following premises:

According to the *predictive coding* hypothesis, state predictions generated during locomotion inference under hamstring muscle vibration should be directionally biased, as during imagined movements, because the prediction process is similar in both. During imagined movements, there is no sensory feedback to be compared with motor prediction, since there is no actual movement. During locomotion inference, however, speed prediction could theoretically be compared with the actor’s actual speed. This would certainly generate an internal error signal, which would be different according to movement direction: locomotion acceleration in the forward direction (remember that imagined locomotion under vibration was faster than normal locomotion, and thus it will be also faster when observing an actor walking forward a natural speed) and locomotion deceleration in the backward direction (imagined locomotion under vibration was slower than normal locomotion, and thus it will be also slower when observing an actor walking backward at a natural speed). Consistent with the *predictive coding* hypothesis, the observer, to provide good estimations about the actor’s walking speed, should try to reduce this error and fit its prediction with the actor’s walking speed. Thus, we expected opposite effects from those found in the first experiment.

On the contrary, if directional effects in the locomotion inference task are similar to those found in the first experiment (i.e., the observer estimates that the actor goes faster during the forward locomotion and vice versa for backward locomotion), we could instead argue for the direct matching hypothesis. Indeed, similar direction effects can only be explained by assuming that observers predict the actor’s movement based on a direct internal simulation of his motion without any internal comparison/correction process. For example, in the forward direction, the observer under vibration feels in a fast walking dynamic, and thus predicts a shorter duration.

ANOVA, applied on the observers’ temporal estimation (plain grey and black histograms, Figure 5), revealed an interaction effect between *vibration* and *direction* (F_(2.16)_=147.1, p<0.0001, η^2^ = 0.95). While vibration of the tibialis anterior muscle did not affect temporal estimation (p>0.1, for both forward and backward locomotion), hamstring muscle vibration induced an overestimation of the actor’s forward walking (p<0.001) and an underestimation of the actor’s backward walking (p<0.001). To further evaluate these vibratory effects (Figure 6), we analyzed the estimation error ([observer’s time estimation – actor’s walking time] / actor’s walking time × 100]). We found a significant effect for forward locomotion (F_(2.16)_=26.8, p<0.0001, η^2^ = 0.77). Precisely, observers accurately estimated the actor’s walking speed in the No-vibration and TA-vibration conditions (p>0.9), but not in the H-vibration (p<0.001). We found comparable results for the backward locomotion (F_(2.16)_=28.1, p<0.0001, η^2^ = 0.78). As for the other experiments, there was not any trial-by-trial adaptation (for all conditions, t<1 and p>0.1, see supplemental Figure 1).

**Figure 5.**
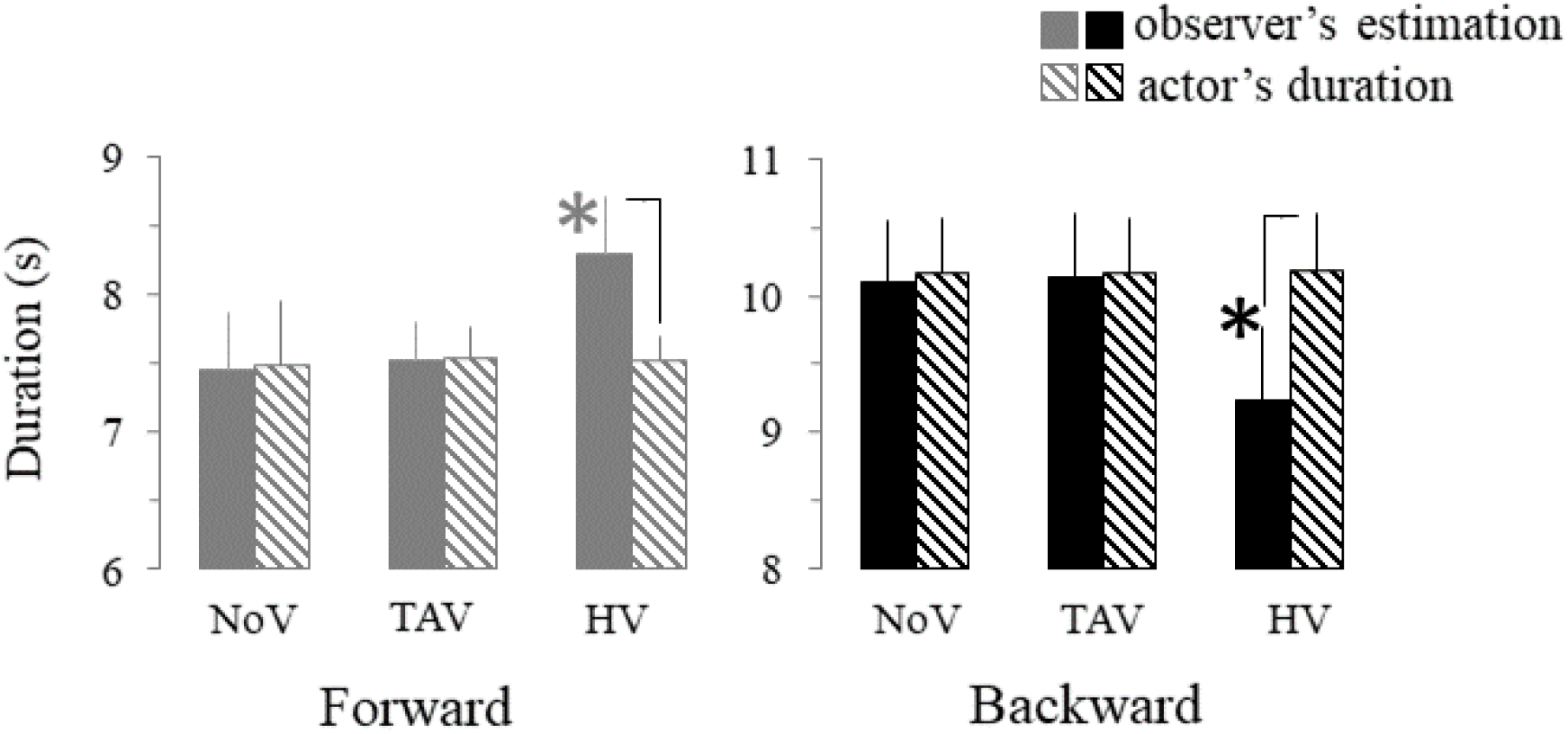
Experiment 3. Forward (grey) and backward (black) locomotion inference for each vibration condition. Results are expressed in terms of mean (+ SD) durations (time in seconds estimated by the observer and performed by the actor on the 9 m linear path). Black lines connecting two bars indicate significant differences between the observer’s estimation and actual actor locomotion duration. Grey and black stars indicate significant differences between HV condition and the two other vibration conditions (NoV and TAV), within the direction of locomotion (forward and backward). NoV, no vibration; TAV, tibialis anterior vibration; H, hamstring vibration.

**Figure 6.**
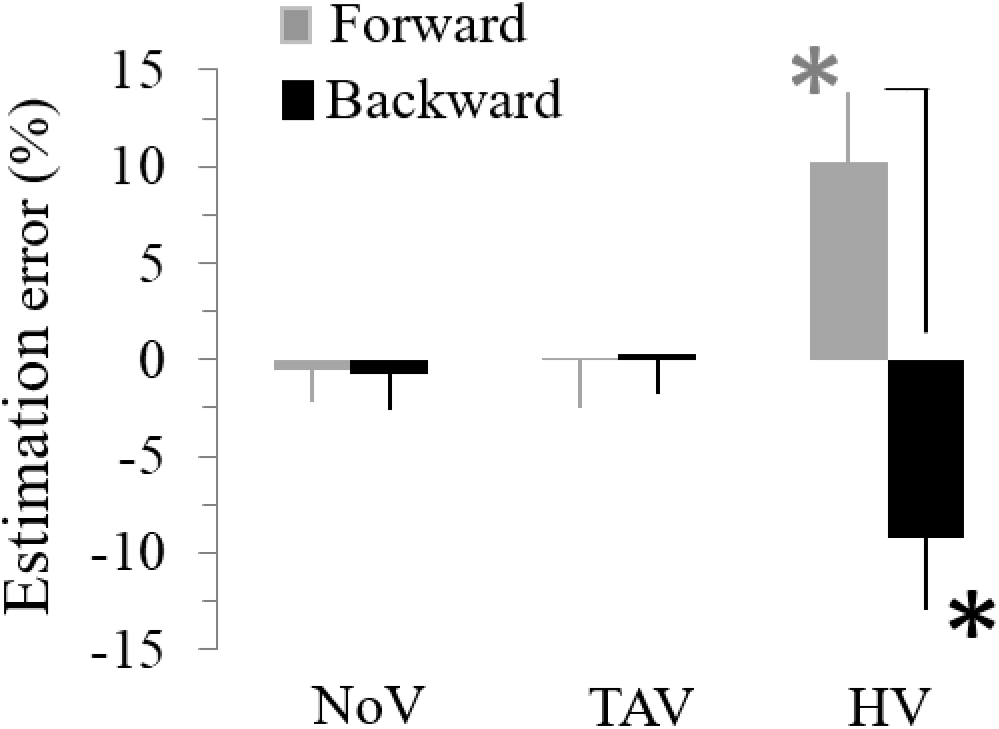
Experiment 3. Mean (+SD) estimation error (observer’s time estimation - actor walking time / observer’s time estimation × 100) for forward (grey) and backward (black) locomotion inference, and each vibration condition. Black lines indicate significant differences between the directions of the walk. Grey and black stars indicate significant differences between HV condition and the two other vibration conditions (NoV and TAV), in forward and backward locomotion, respectively. NoV, no vibration; TAV, tibialis anterior vibration; H, hamstring vibration.

Note that hamstring muscle vibration at 40 Hz (control experiment 3a, see supplemental Table 1) did not alter the temporal estimation of the actor’s speed in forward (Permutation test; t=0.04, P=0.96) and backward locomotion (Permutation test; t=-0.16, P=0.84).

After the locomotion inference experiment, we questioned participants on the strategy they used to estimate the actor’s walking speed. All reported mentally simulating the actor’s movement from their body perspective (first-person perspective). To test whether another strategy, such as external visual imagery (third-person perspective), that does not involve motor prediction, would produce similar results, we explicitly requested six participants that participated in Experiment 3, to repeat the experiment by using visual imagery to estimate actor’s locomotion speed (control experiment 3b). The actor’s forward walking durations and the corresponding estimated durations of the observers were not different (in all comparisons, t<1 and p>0.1; see supplemental Table 2 for mean durations and standard deviation). Estimation errors were very low (individual values ranged between −1.13% to 1.10%) and not different between conditions (p>0.5).

### Comparison between locomotion inference and mental locomotion

Interestingly, hamstring muscle vibration induced almost opposite effects on locomotion inference and mental locomotion (Figure 7). Indeed, the statistical comparison of the vibratory effects, assessed by the ratio [(HV – NoV) / NoV*100)] and calculated for locomotion inference and mental locomotion separately, showed a significant effect for the forward (t=-9.84, p<0.0001) and backward locomotion (t=13.82, p<0.0001).

**Figure 7.**
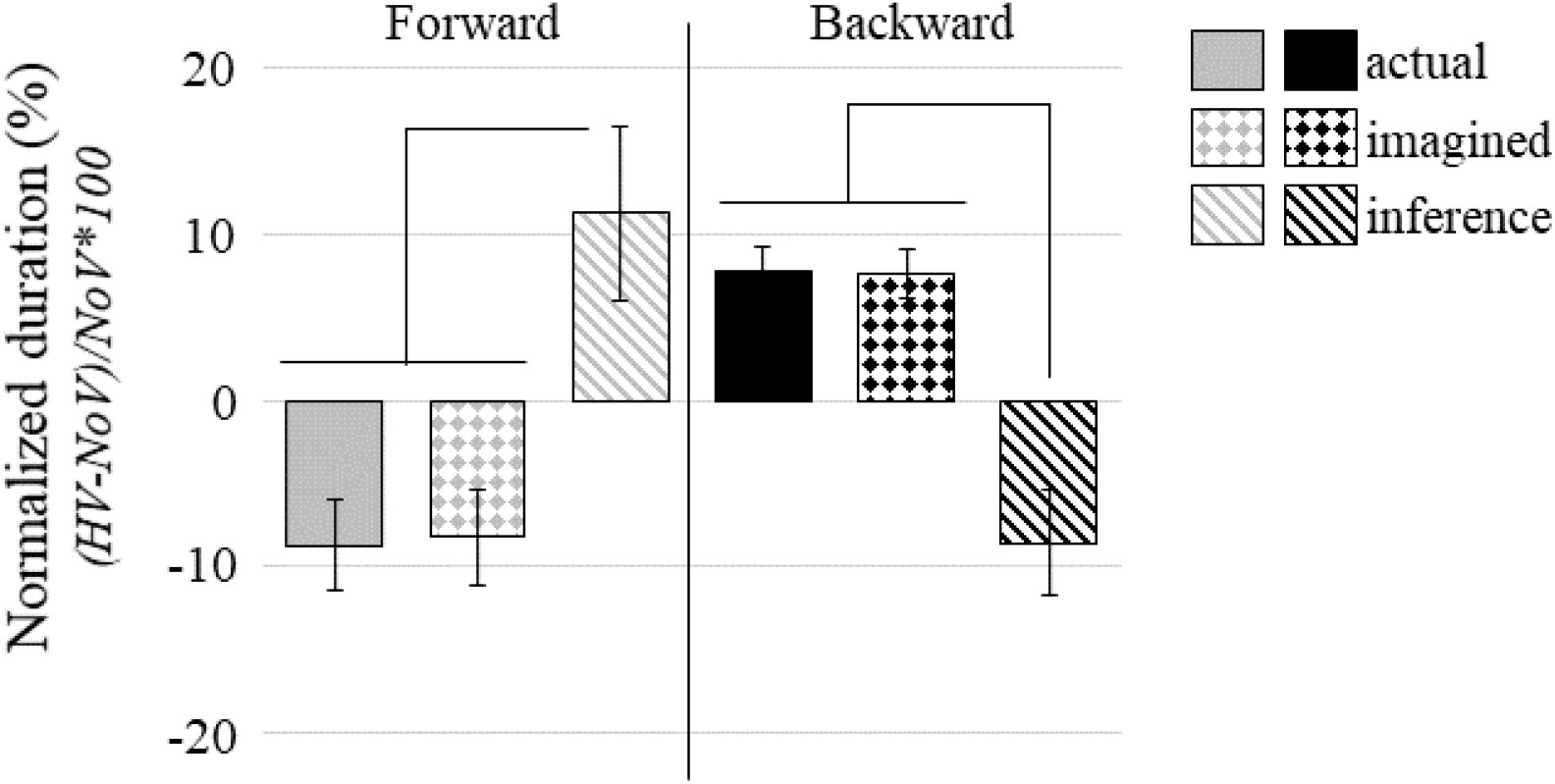
Effects of hamstring muscle vibration (normalized by no vibration, NoV), on actual locomotion, imagined locomotion, and locomotion inference, for forward and backward walking. Black lines connecting two bars indicate significant differences between the conditions within each direction of locomotion.

### Experiment 4: Motor inference of locomotion under different conditions of visual feedback

In this experiment, we hypothesize that the amount of visual feedback available will have a significant impact on motor prediction and thus on the observers’ ability to accurately estimate the duration of the actor’s locomotion. We asked six new participants (2 females, 4 males, mean age: 23.9 ± 2.2 years) to perform the same experiment with some modifications in the experimental design. The actor walked only forward and only the hamstring muscle of the observers was vibrated.

In the first condition (similar to experiment 3), the actor adopted the natural walking speed of each observer, who perceived the actor’s locomotion for 1/3 of the path without being vibrated. In the second condition (new), the actor adopted a higher speed than the natural walking speed of each observer, who perceived the actor’s locomotion for 1/3 of the path without being vibrated. In the third condition (similar to experiment 3), the actor adopted the natural walking speed of each observer, who perceived the actor’s locomotion for 1/3 of the path while being vibrated. In the fourth condition (new), the actor adopted the *‘estimated’* walking speed of each observer; that is the actor walked at the speed that the observer indicated while being vibrated (see previous condition). The observer perceived the actor’s locomotion for 1/3 of the path while being vibrated. In the fifth condition (new), the actor adopted the natural walking speed of each observer, who perceived the actor’s locomotion for 2/3 of the path while being vibrated. Each participant realized 12 trials in each condition. The actor’s and observers’ duration were recorded as described in Experiment 3.

As required by the task’s instructions, the actor increased the walking speed in the ‘higher speed’ and ‘matched speed’ conditions (Figure 8). The permutation tests ran on the actor’s forward walking durations showed that these two conditions were different from the 3 ‘natural speed’ conditions (all p<0.05). We found no significant difference between the actor’s forward walking durations in the ‘higher speed’ compared to ‘matched speed’ conditions (p=0.52, t=-1.63). These results confirmed that the task instructions were respected by the actor.

**Figure 8.**
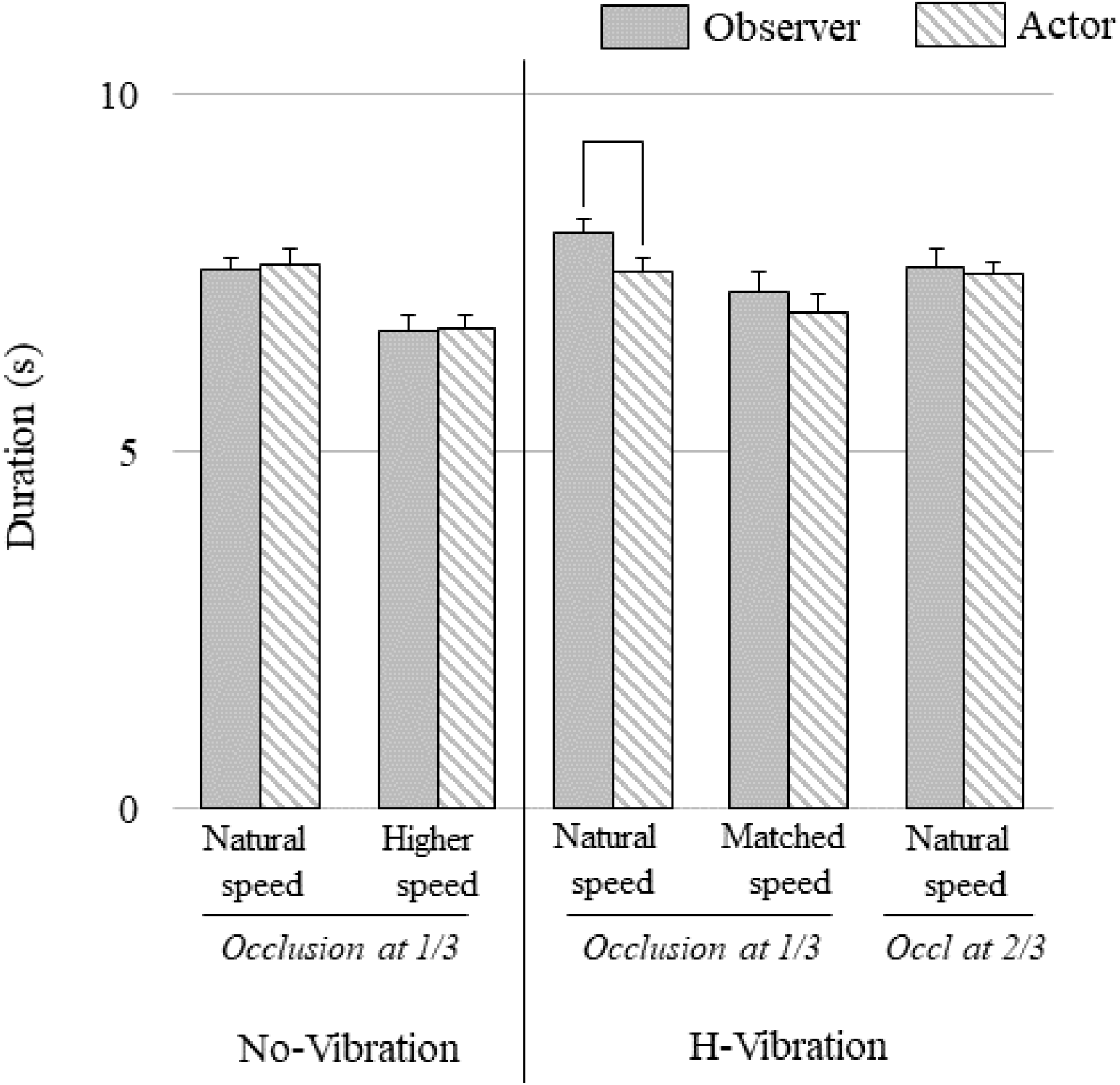
Experiment 4. Mean (+ SD) duration of the actor’s locomotion (grey striped bars) and mean (+ SD) durations estimated by the observer (plain grey bars), for the five experimental conditions of the forward locomotion inference task. In conditions 1 and 2 no vibration (NoV) was applied to the observer, while in conditions 3 to 5 a hamstring muscle vibration (HV) was applied. In the ‘Natural speed’ conditions, the actor walked at the observer’s natural speed. In the ‘Higher speed’ condition, the actor walked at a higher speed than the natural one (+15%). In the ‘Matched speed’ condition, the actor walked at the speed of the observer under HV. Finally, the time at which the visual occlusion appeared for the observer was manipulated (occluded after 1/3 of the movement for conditions 1 to 4, and after 2/3 of the movement for condition 5). Black lines connecting two bars indicate significant differences between the real duration of the walk (actor’s walk duration) and the duration estimated by the observer.

For the observers, we replicated the effects of Experiment 3 for the same conditions. The observers accurately estimated the actor’s walking speed without vibration (condition 1: ‘Natural speed, 1/3’ without vibration; p=0.95, t=-0.52), white they overestimated it under vibration (condition 3: ‘Natural speed, 1/3’ under H-vibration; p<0.01; t=5.31).

For condition 2 (Higher speed, 1/3, without vibration), we found that when the actor increased its walking speed, the observer still accurately estimated it (p=0.97, t=-0.47) suggesting that locomotion inference was speed-independent. More interestingly, in condition 4 (Matched speed, 1/3’, vibration), when the actor’s speed matched that of the observers under hamstring muscle vibration (that is the actor was walking faster), their estimation was accurate (no underestimation; p=0.31, t=1.90). Compared to the same condition at natural speed (condition 3), in which the observers underestimated the actors’ speed, this novel finding indicates that the initial mismatch between the observers’ speed prediction and the actor’s speed constitutes an internal error signal that regulates speed estimation during locomotion inference. Finally, in condition 5 (Natural speed, 2/3, vibration), speed underestimation disappears (p=0.75, t=0.99) when the observer was able to perceive for a longer time the actor’s movement (6 meters). In line with the *predictive coding* hypothesis, the greater amount of feedback available here may allow the observer to adjust the quality of his prediction to match the actor’s actual walking speed as closely as possible.

## Methods

### Participants

All participants gave written informed consent to participate in the experiments, which were approved by the local Ethics committee of the *Université de Bourgogne* and conducted following the Declaration of Helsinki. Participants were free from muscular, cognitive, or nervous disorders and were sensitive to vibratory stimulation from the first application. None of them had previous experience concerning the sensorimotor effects caused by musculotendinous vibration and were unaware of the specific hypotheses being tested in the study.

### Experiment 1: Imagined locomotion under muscle vibration

Nine volunteers (5 females, 4 males, mean age: 25.4 ± 2.7 years) participated in this experiment, which took place in a quiet and large room (10 × 20 m), temperature regulated (22 ± 2 °C), and illuminated with homogeneous white light. The ground was smooth and plain. On the floor, two black lines materialized the initial and the final position of a walking path (9m). Participants were standing upright behind the starting line, their arms were hanging along the body, and their feet were parallel and slightly apart (Figure 1). They had to mentally simulate (motor imagery) walking at a natural speed along the path in a forward or backward direction, with eyes open to facilitate the visualization of the path, and under three vibration conditions: No-Vibration (NoV), bilateral vibration of the tibialis anterior muscles (TAV), and bilateral vibration of the hamstring muscles (HV). We instructed participants to feel themselves performing the motor task in a first-person perspective (i.e., motor or kinesthetic imagery, as in Rannaud Monany et al., 2022; Truong et al., 2022), rather than just watching themselves doing it (visual imagery). The participants realized 12 trials in each experimental condition (n= 72; 12 trials × 2 walking directions × 3 vibrations). The conditions of vibration (NoV, TAV, and HV) were spaced out by at least twenty-four hours, to avoid interferences due to the post-effects of muscle vibration (Courtine et al., 2001; Wierzbicka et al., 1998) and were counterbalanced among the participants. Within each vibration condition, forward and backward locomotion was also counterbalanced among the participants. The duration of mental movements was recorded using an electronic stopwatch (temporal resolution 1 ms) that participants hold in their left arm. They started the stopwatch when they mentally started to walk and stopped it when they mentally reached the end line. This mental chronometry paradigm is known to provide reliable results (for example Demougeot & Papaxanthis, 2011; Gueugneau et al., 2008, 2015). The experimenter only had access to the recorded durations so participants had no information about their imagined walking speed. Mechanical vibration began 5 s before the onset of the imagined movement and ended 5s after its achievement. All the participants verbally reported at the end of the experiment that they imagined walking in a first-person perspective without difficulties. Additionally, after each block of imagined trials, the participants reported the subjective estimation (SE) of the imagined movement quality through a 7-point Likert scale (1: very hard to feel and see the movement, 7: very easy to feel and see the movement, 2–6: intermediate score).

### Control Experiment 1a

To control the effects of vibration intensity on mental locomotion, Experiment 1 was replicated by setting this time the muscles’ vibration at 40 Hz. Five participants that already participated in Experiment 1 (2 females, 3 males, mean age: 24.4 ± 2.1 years), took part in this control. This control was always realized two days after Experiment 1.

### Control Experiment 1b

Muscle vibration of the tibialis anterior and hamstring muscles caused postural reactions (Figure 1). To control, whether these postures could affect mental walking durations, we asked five participants, who already participated in Experiment 1 (1 female, 4 males, mean age: 23.9 ±1.8 years) to imagine forward walking without any muscle vibration, but by adopting the postures elicited by the vibration of the tibialis anterior and hamstring muscles. This control was always realized three days after Experiment 1.

### Experiment 2: Actual locomotion under muscle vibration

The nine participants of Experiment 1 took also part in Experiment 2, at least one week after their involvement in Experiment 1 and the control Experiments 1a and 1b. The experimental design was identical to that of Experiment 1 (i.e., 2 walking directions × 3 vibrations) and was realized in the same environment. Participants were requested to walk along the 9 m linear path, at a natural speed, with full vision. Walking along the path at self-paced velocity required at least five gait cycles. Participants accomplished 12 trials for each experimental condition. The walking duration was recorded using an electronic chronometer. Participants started the stopwatch when they started to walk and stopped it when they reached the end line. As in Experiment 1, the experimenter only had access to the recorded durations so participants had no information about their imagined walking speed. The mechanical vibration began 5 s before the onset of the locomotion and ended 5s after its achievement.

### Control experiment 2

To control the effects of vibration intensity on actual locomotion, Experiment 2 was replicated by setting this time the muscle vibration at 40 Hz. Five participants who participated in Experiment 2 (2 females, 3 males, mean age: 23.1± 3.0 years) took part in this control.

### Experiment 3: Locomotion inference under muscle vibration

A new group of nine volunteers (4 females, 5 males, mean age: 25.2 ± 2.1 years), novices regarding the effects of muscle vibration on actual and mental locomotion, participated in Experiment 3, which took place in the same environment as the two others. Each participant (hereafter ‘observer’) had to estimate the duration needed by another human (hereafter ‘actor’) to walk along the same 9 m linear path. The actor walked forward or backward, with full vision, at natural velocity, without muscle vibration. A preliminary practice allowed the actor to adapt its speed to the natural speed of each observer with small variability. This was possible because, before the start of the experiment, we asked the observers to naturally walk along the path (ten trials), while measuring the duration of their locomotion. The observer stood upright in front of the actor at a distance of 2 m. The observer could fully perceive the actor’s locomotion for 1/3 of the path (3 m). For the rest of the path (6 m), the vision of the observer was occluded by crystal goggles with a wireless obstruction system (*Plato System*, 2 ms resolution). The actor’s walking duration and the corresponding temporal estimation by the observer were recorded using two electronic chronometers manipulated by the actor and the observer and synchronized via a computer program (see more details below). The observer realized the task under the same vibration conditions (NoV, TAV, and HV) as in the previous experiments. Vibrations were spaced out by at least twenty-four hours, to avoid interferences due to the post-effects of muscle vibration) and were counterbalanced among the participants. Within each vibration condition, forward and backward locomotion was also counterbalanced among the participants. The experimenter only had access to the recorded durations so participants had no information about their temporal estimation and mechanical vibration began 5 s before the onset of the actors’ locomotion and ended 5s after its achievement.

### Control experiment 3a

To control the effects of vibration intensity on locomotion inference, Experiment 3 was replicated by setting this time the muscle vibration at 40 Hz. Five new participants (1 female, 4 males, mean age: 21.8± 2.9 years) took part in this control.

### Control experiment 3b

By the end of the locomotion inference experiment, we questioned participants on the strategy they used to estimate the actor’s walking duration. All reported mentally simulating the actor’s movement from their body perspective, i.e., first-person perspective (see above motor imagery). To test whether another strategy, such as external visual imagery (i.e., third-person perspective), would produce similar results, we explicitly asked six participants (2 females, 4 males, mean age: 24.6 ± 1.9 years) who participated in Experiment 3 to reproduce the experiment by visualizing this time the actor’s locomotion. All participants reported generating clear visual images in both directions and under all vibration conditions. The experimental design was strictly similar to Experiment 3.

### Experiment 4: Motor inference of locomotion under different conditions of visual feedback

To further explore the mechanisms underlying locomotion inference, we asked six new observers (2 females and 4 males, mean age: 23.9 ± 2.2 years) to perform the same experiment with some modifications in the experimental design. Among the five following conditions, the 1^st^ and 3^rd^ were similar between Experiments 3 and 4, while the other three were new.

In experiment 4, the actor walked only forward and only the observers’ hamstring muscles were vibrated. All the conditions were realized on different days and were counterbalanced among the participants. Hamstring muscle vibration was similar to the previous experiments and only the experimenter had access to the recorded durations. In the first condition (NoV, Natural speed, 1/3 of the path), the actor adopted the natural walking speed of each observer, who perceived the actor’s locomotion for 1/3 of the path without being vibrated. In the second condition (NoV, Higher speed, 1/3), the actor adopted a higher speed (+15%) than the natural walking speed of each observer, who perceived the actor’s locomotion for 1/3 of the path without being vibrated. In the third condition (HV, Natural speed, 1/3 of the path), the actor adopted the natural walking speed of each observer, who perceived the actor’s locomotion for 1/3 of the path while being vibrated. In the fourth condition (HV, Matched speed, 1/3), the actor adopted the *‘estimated’* walking speed of each observer; that is the actor walked at the speed that the observer indicated while being vibrated (see previous condition). The observer perceived the actor’s locomotion for 1/3 of the path while being vibrated. In the fifth condition (HV, Natural speed, 2/3), the actor adopted the natural walking speed of each observer, who perceived the actor’s locomotion for 2/3 of the path while being vibrated. Each participant realized 12 trials in each condition. The actor’s and observers’ duration were recorded as described in Experiment 3.

### Movement duration recording

In all experiments, the duration of mental, actual, and inferred locomotion was recorded using an electronic stopwatch (temporal resolution 1 ms) that participants hold in their left arm. The specific procedure is explained in each experiment.

### Muscle vibration

Tibialis Anterior and Hamstring muscles were bilaterally stimulated by two mechanical vibrators (VB115, Technoconcept, France, biaxial DC motors, 7cm long, with a radius of 4.5 cm and a weight of 150 g, delivering a 1.5 mm amplitude/80 Hz sinusoid signal). Vibrators were cautiously fastened to the participants’ limbs with elastic bands in a way that did not disturb natural locomotion or posture. The tension developed by the elastic bands (3 cm wide) was about 7 N cm^−1^, which corresponded to a pressure of about 1 N cm^−2^ (i.e., 75 mmHg acting on the arterial vessel walls, which is about 50% of the compression value necessary to occlude them while in standing position). The vibrators were fixed on the Tibialis Anterior (~4 cm above the ankle joint) and the Hamstring distal (~3 cm above the knee joint) tendons.

These two muscles were selected after a pre-test session ran on five participants, who did not take part in the main experiments. We tested the effects of Triceps Surae, Tibialis Anterior, Hamstring, and Splenius muscle vibration on body posture and walking speed. Following previous studies (Eklund, 1972; Ivanenko et al., 2000; Courtine et al., 2007), we found that vibration of Triceps Surae and Tibialis Anterior muscles had a strong postural effect, namely a backward and a forward body tilt, respectively, but no effect on walking speed. On the other hand, the vibration of the Hamstring and Splenius muscles had both postural and walking effects. Hamstring muscle vibration elicited backward inclination of the trunk and forward inclination of the shank due to hip extension and knee flexion. Splenius muscle vibration induced a forward body tilt. In addition, both vibratory stimulations accelerated forward locomotion and decelerated backward locomotion. According to these preliminary results, and to alleviate the whole protocol, we decided to vibrate the Tibialis Anterior and Hamstring muscles during the main and control experiments.

### Electromyographic activity

We recorded the EMG activity (SMART system, BTS, Italy) of the lower limb muscles during experiments 1 (Mental locomotion) and 3 (Locomotion inference), as well as during 12 trials of rest in which participants were erect and immobile for 8 s without performing any task. EMG activity was recorded using two silver-chloride bipolar surface electrodes from eight muscles belly: soleus, gastrocnemius medialis, gastrocnemius lateralis, tibialis anterior, biceps femoris, rectus femoris, vastus medialis, and vastus lateralis. A ground electrode was placed on the forehead. EMG signal was pre-amplified, digitized, and transmitted to the remote amplifier via an optic fiber. It was sampled at 1000 Hz, filtered between 20-400 Hz (Butterworth), and rectified for further analysis. To quantify muscle activation, we calculated the root mean square (RMS) values of the EMG signal, using the following formula:

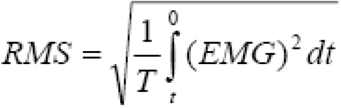

RMS values during all trials ranged from 2.1 mV and 5.65 mV and were not significantly different from those of the rest trials (*two-tailed paired t-tests, p*>0.05, for all comparisons).

### Statistical analysis

We computed individual average durations and standard deviations (SD) for each experimental condition and we verified that all variables were normally distributed (Kolmogorov–Smirnoff test) and that sphericity was respected. An a priori G*POWER analysis for total sample size estimation (parameters: Effect size f = 0.42; α = 0.05; power = 0.90; partial η^2^ = 0.15; groups = 1; number of measurements = 6), indicated 9 participants. The data collected were processed with STATISTICA (8.0 version; Stat-Soft, Tulsa, OK) and custom software written in Matlab (Mathworks, Natick, MA).

For Experiments 1, 2, and 3, we tested the effect of *vibration* (NoV, TAV, and HV) and *direction* (Forward, Backward) by performing two-way repeated measurements ANOVA, with Scheffé *post-hoc* comparisons when necessary. To support our results, eta-squared (η^2^) effect sizes were calculated for the ANOVAs.

To compare the effects of muscle vibration across the experiments, we normalized the durations obtained in the muscle vibration conditions (H-V or TA-V) by the durations obtained in the No-Vibration condition (NoV): condition (V-NoV)/NoV*100. To compare the magnitude of the effects, we then performed two-way (Vibration × Direction) repeated measurements ANOVA. In addition, to further evaluate the effect of muscle vibration on mental and actual locomotion, we performed Pearson’s correlation on the normalized values between the conditions.

Finally, to test the adaptation effects within the 12 trials of Experiments 1, 2, and 3, we averaged the first three trials (1-3) and the last three trials (10-12) for each participant and compared them using two-tailed *t-tests* for dependent samples.

For Experiment 4 and control experiments, data were not normally distributed. We performed multiple permutation tests using the Matlab function ‘mult_comp_perm_t2’ using 5000 repetitions. When applying permutation tests with multiple comparisons a correction must be performed. The “tmax” method was used for adjusting the p-values of each variable in the same way as Bonferroni correction does for a t-test (Blair & Karniski, 1993; Westfall & Young, 1993).

## Discussion

In the present study, we experimentally evaluated the *direct matching* and the *predictive coding* hypotheses, candidates both to explain the underlying functional process during action observation. The first assumes a direct equivalence between the visual information of the actor’s movement and the activation of the corresponding motor representations in the observer’s motor cortex. The second postulates that motor predictions are generated and compared to the visual information of the actor’s movement. To discriminate between them, we used a locomotor task under different experimental conditions. We asked participants either to imagine walking, a well-known paradigm to measure motor prediction (Courtine et al., 2004; Personnier et al., 2010), or to watch an actor walking and to evaluate the moment at which he/she would reach the end of the path (inference locomotion, Pozzo et al., 2006; Saunier et al., 2008). In addition, we manipulated imagined and inference locomotion by creating sensory illusions via peripheral mechanical muscle vibration. We found that muscle vibration accelerated imagined forward locomotion and decelerated imagined backward locomotion, compared to imagined locomotion in the non-vibrated condition. Similar results were obtained for actual walking. Intriguingly, we found opposite results for locomotion inference: observers estimated longer and shorter durations for an actor walking forward and backward, respectively. Such findings argue in favor of a common sensorimotor prediction process involved in action observation and action mental simulation.

An important feature of motor behavior is the ability to predict the consequences of actions before sensory feedback is available. Evidence supports the hypothesis that motor prediction is generated by internal forward models, which are neural networks that mimic the causal flow of the physical process by predicting the future sensorimotor state (e.g., position, velocity) given the efferent copy of the motor command and the current state (Miall & Wolpert, 1996; Wolpert & Flanagan, 2001). Motor prediction is useful in motor learning through mental practice (Guillot et al., 2021; Ruffino et al., 2017a, 2017b, 2021) It is proposed that the temporal features of mental movements emerge from sensorimotor predictions of the forward model (Papaxanthis et al., 2012; Sirigu et al., 1996). Here, we showed that sensory illusions evoked by muscle vibration had specific directional effects on the timing of imagined locomotion: with respect to the non-vibration condition; it accelerated forward and decelerated backward locomotion. This is possible because, during motor imagery, we internally simulate/predict upcoming sensory experiences (position, speed) based on the prepared but blocked motor commands. It is of interest that similar results are obtained during actual locomotion. This suggests that sensory illusions and the consistent locomotor effects under muscle vibration are the consequence of central mechanisms operating in both actual and imagined movements and not the outcome of unspecific effects, such as an extra load that would affect attention or perturbed movement execution.

The results of control experiments 1a and 1b also validated this conclusion. In line with this, it has been shown that motor imagery and the illusory sensation during arm movements commonly activated the contralateral cingulate motor areas, supplementary motor area, dorsal premotor cortex, and ipsilateral cerebellum. In addition, previous studies have proposed that the combination of a proprioceptive evoked sensation with an imagined movement generates movements that were the sum of the vectors of the sensations evoked by the two modalities (Kitada et al., 2002; Naito et al., 2002b; Shibata et al., 2017; Thyrion & Roll, 2009).

We found that sensory illusions elicited by muscle vibration had also a specific directional effect during locomotion inference. This effect was, however, opposite to that observed during imagined locomotion: observers estimated longer and shorter durations for an actor walking forward and backward, respectively. We consider that this pattern of results is in line with the *predictive coding* hypothesis. It denotes that the observer estimates the actor’s speed, not by purely reproducing the actor’s movement but by comparing the observed action to his/her prediction. When observing forward locomotion under muscle vibration, the observer is in a sensory predictive state that accelerates walking speed (see imagined locomotion). Therefore, when comparing the actor’s speed to his/her speed prediction, the observer estimates that the actor is going slower and infers a longer overall duration compared to the condition without vibration. The opposite reasoning can be made for backward locomotion. Note that these effects were due to the sensory effects of muscle vibration at 85Hz. Vibration at 40Hz did not cause any sensory effect and thus any bias in the observers’ estimation.

This directional effect suggests that to estimate actors’ motion, the observer uses an internal error signal, an issue from the comparison of the actors’ walking speed and his/her speed prediction. Note that the observers can provide accurate estimations of the actor’s locomotion in the non-vibration condition, no matter if the actor is walking at a natural or high speed (experiment 4), suggesting that the previous findings are due to sensorimotor prediction/comparison rather than to the observes’ ability to accurately estimate speed locomotion. In addition, when the prediction error was annulated, by asking the actor to match the predictive speed of the observers under muscle vibration (experiment 4), the observer accurately estimated the actors’ speed. This further indicates that it is a predictive/comparison process that is used for locomotion inference. Finally, sustained visual feedback (2/3 of the path, experiment 4) can provide valuable information to the observer to adapt its estimation for the actor’s speed.

The *predictive coding* hypothesis has been enriched by several theoretical papers (Friston et al., 2011b; Keysers & Gazzola, 2014; Keysers & Perrett, 2004; Kilner & Frith, 2008). Recent neurophysiological evidence seems also to support it. Hilt and colleagues (Hilt et al., 2020) found a significant correlation between the level of cortico-spinal excitability (TMS on M1) and the degree of similarity between the observer’s own motor style and the observed motor style. Interestingly, the correlation they found was negative, i.e., the greater the degree of similarity, the smaller the associated MEPs. This increase in MEP amplitude may reflect the error-predictive signal from the cerebellum to the motor cortex. It is of great interest that previous studies have shown stronger and longer effects on neuroplasticity when action observation is combined with peripheral stimulation (Bove et al., 2009; Bisio et al., 2017; Bisio, Avanzino, Gueugneau, et al., 2015). For example, vibratory muscle stimulation in conjunction with action observation has also shown a sustained increase in activity in M1; up to 1 hr after stimulation (Bisio et al., 2019). The authors suggested that during action observation the brain exploiting the visual input channel uses information from multiple senses to represent the body and act on the world.

According to the *directing matching* hypothesis, two results, different from those described above, could be expected. If one assumes that the observers’ sensorimotor state is not included in the action observation process, then: (i) sensory illusions should not have any effect on locomotor inference. This was not obviously the case. However, when observers were explicitly requested to use visual imagery (third-person perspective) to estimate the actor’s movement duration, we found no difference between actual and estimated duration. This suggests that although the use of purely visual information during locomotion inference is possible, the authors did not spontaneously use this strategy. (ii) similar effects to imagined movements should be obtained. As the observer under muscle vibration feels walking faster, he/she should estimate a shorter duration for the actor’s movement.

In conclusion, this study goes further in understanding the mechanisms underlying action observation. We have shown that an internal prediction process, including a comparison between the observer’s state prediction and the external visual feedback, is involved in action observation. This makes action observation a dynamic process during which internal prediction is updated until it matches external visual information.

## Notes

### Competing Interest Statement

The authors have declared no competing interest.

