## Supplementary Material for "Evidence for a sensorimotor prediction process in action observation"

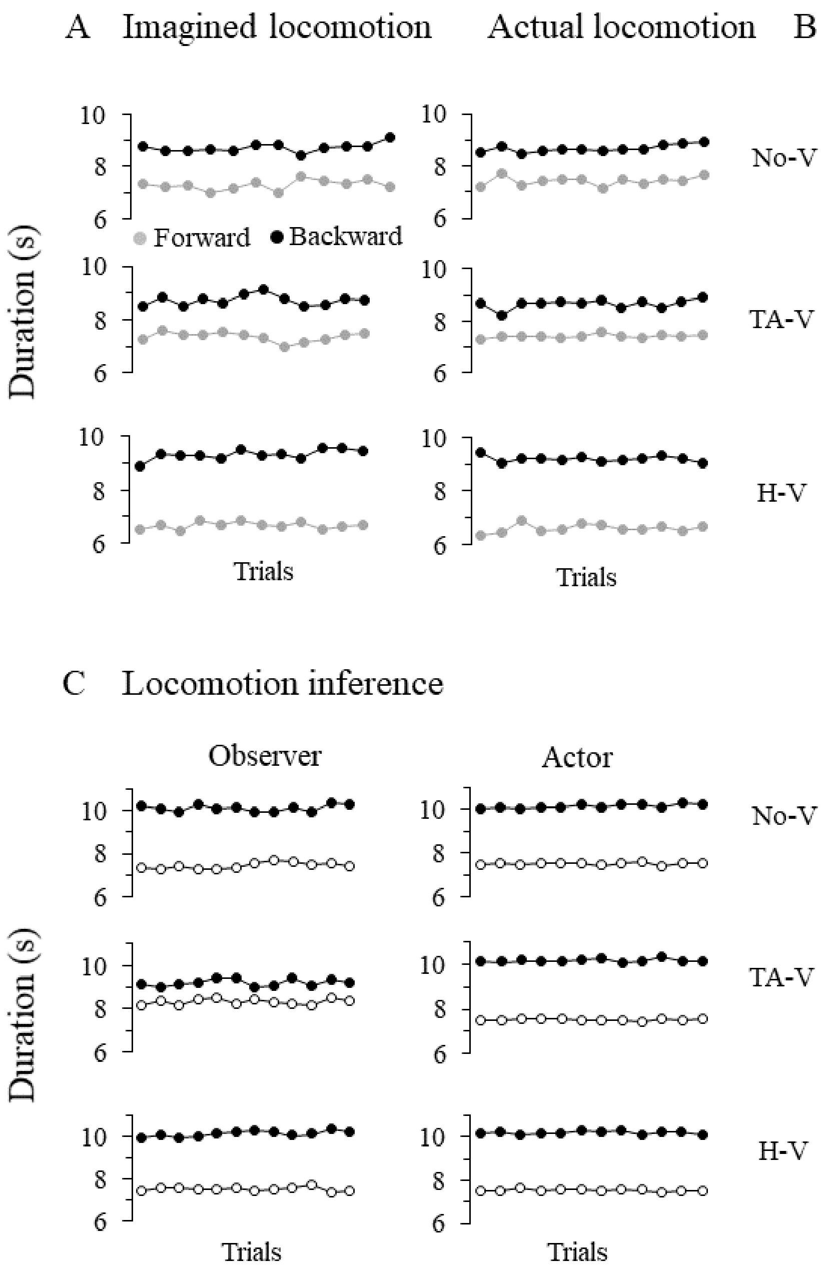


Figure 1: Mean trial-by-trial durations (in seconds) for forward (grey) and backward (black) imagined walking (A, Experiment 1) and actual walking (B, Experiment 2), in the three vibration conditions. (C) Mean trial-by-trial temporal estimations of the observers, and trial-by-trial mean movement durations of the actor during forward (white) and backward (black) walking, in the three vibration conditions. To statistically evaluate the stability across trials in each condition, we compared the mean over the first three trials (1-3) and over the last three trials (10-12) and showed no significant difference (all $t<1$ and $p>0.1$). NoV, no vibration; TAV, tibialis anterior muscle vibration; HV, hamstring muscle vibration.

Table 1: Mean durations (±SD) of forward and backward locomotion for the control experiments 1a (imagined locomotion with or without muscular vibration at 40Hz), 1b (imagined locomotion in different starting postures), 2 (actual locomotion with or without muscular vibration at 40Hz) and 3a (locomotion inference with or without muscular vibration at 40Hz).

| Duration (s) | Forward | Backward |
|---|---|---|
| <u>Control experiment 1a</u> | | |
| No-vibration imagined locomotion | 7.14 ±0.52 | 8.30 ±0.49 |
| H-vibration imagined locomotion | 7.12 ±0.49 | 8.32 ±0.52 |
| <u>Control Experiment 1b</u> | | |
| Erect Posture | 7.01 ±0.55 | |
| Posture corresponded to TA-vibration | 7.06 ±0.63 | |
| Posture corresponded to H-vibration | 6.99 ±0.54 | |
| <u>Control experiment 2</u> | | |
| No-vibration actual locomotion | 7.28 ±0.59 | 8.45 ±0.53 |
| H-vibration actual locomotion | 7.31 ±0.60 | 8.50 ±0.55 |
| <u>Control experiment 3a</u> | | |
| No-vibration actor | 7.41 ±0.22 | 10.24 ±0.24 |
| H-vibration observer (40Hz) | 7.41 ±0.24 | 10.21 ±0.22 |

Table 2: Mean durations (±SD) in the locomotion visualization task (control experiment 3b). Statistical comparions between actor’s forward walking durations and the corresponding estimated durations showed no significant difference (all $t<1$ and $p>0.1$).

| Duration (s) | Forward | Backward |
|---|---|---|
| No-vibration | | |
| Observer’s estimation | 7.40 ±0.24 | 10.23 ±0.21 |
| Actor’s duration | 7.41 ±0.22 | 10.24 ±0.24 |
| H-vibration | | |
| Observer’s estimation | 7.39 ±0.23 | 10.23 ±0.21 |
| Actor’s duration | 7.36 ±0.21 | 10.26 ±0.21 |
